# Identification of potential biomarkers in subtypes of epithelial ovarian cancer

**DOI:** 10.1101/2020.02.24.962472

**Authors:** Rinki Singh, Anup Som

## Abstract

Epithelial ovarian cancer (EOC) is the most lethal gynecological cancer. Due to the lack of specific symptoms, ∼80% of epithelial ovarian cancer is diagnosed at an advanced stage and often metastasize to the distant organ. Epithelial ovarian cancer is a heterogeneous disease that is classified into four major histological subtypes namely, serous carcinoma (SC), endometrioid carcinoma (EC), mucinous carcinoma (MC), and clear cell carcinoma (CCC). Ovarian cancer treatment is complicated due to the heterogeneity of the tumors. Patients with different subtypes respond differently to the same treatment and also have different prognoses. This diversity extends to various clinical outcomes of the disease. Thus, identifying new reliable potential biomarkers irrespective of their subtypes is an urgent need for the diagnosis and prognosis of epithelial ovarian cancer. In this study, we performed comparative gene expression analysis for identifying potential biomarkers in four histological subtypes of epithelial ovarian cancer (EOC) that include serous, endometrioid, mucinous, and clear cell carcinomas. Differentially expressed genes (DEGs) between cancerous and normal tissue samples were identified by considering the criteria of absolute logarithmic fold change |log_2_fc|>1 and adjusted p (p_adj_) value<0.05. Pathway enrichment analysis of the DEGs showed that pathways in cancer, PI3K-AKT signaling pathway, RAP1 signaling pathway, cell cycle, cell adhesion molecules, and proteoglycans in cancer were common among the selected cancer subtypes. Further, we constructed the co-expression network of DEGs and identified 15 candidate genes. Finally, based on the survival analysis of the candidate genes, a total of nine genes namely ASPM, CDCA8, CENPM, CEP55, HMMR, RACGAP1, TPX2, UBE2C, and ZWINT with significant prognostic value was proposed as the potential biomarker.

## 1. INTRODUCTION

Ovarian cancer (OC) is one of the most common gynecological cancers that have the worst prognosis and the highest mortality rate in women worldwide (Momenimovahed et al., 2019). The cause of the high mortality rate is advance stage diagnosis, which is metastasizing in the peritoneal cavity. Early detection of ovarian cancer is difficult because of its asymptomatic growth and lack of reliable screening techniques and chemoresistance of recurrent diseases. Previous studies showed that up to 90% of all OCs have an epithelial origin (Torre et al., 2018). Epithelial ovarian cancer (EOC) is further classified into four histologic subtypes namely serous, mucinous, endometrioid, and clear cell carcinomas. Each of these subtypes is associated with different genetic risk factors and molecular events during tumor growth and characterized by distinct mRNA expression profile due to this each subtype is treated as different disease (Kurian et al., 2005; Prat, 2012). High-grade serous and endometrioid subtypes were associated with better survival, whereas patients with mucinous and clear cell subtypes were associated with a higher risk of mortality (Zhou et al., 2018). Carbohydrate antigen 125 (CA125) is the most robust and well-known serum biomarker used for the detection of ovarian cancer. However, it is reported that CA125 is highly expressed in the serous subtype than endometrioid, clear and often normal in mucinous carcinoma. Thus, CA125 not be a highly specific diagnostic marker for ovarian cancer (Choi et al., 2018). The standard treatment for ovarian cancer is cytoreductive surgery and platinum-based chemotherapy, along with the target-based therapy is also used. Ovarian cancer treatment is complicated by the heterogeneity of tumor. Patients with different subtypes respond differently to the same treatment and also have different prognoses (Kim et al., 2018). Thus, identifying new reliable potential biomarkers irrespective their subtypes is an urgent need for the diagnosis and prognosis of the epithelial ovarian cancer.

Over the past decades, various finding and focused research has been carried out to understand the underlying molecular mechanism of the ovarian cancer and its diagnosis (Hibbs, 2004; Cai et al., 2015; Krzystyniak et al., 2016; Willis et al., 2016, Singh and Som, 2019). Moreover, a single gene-based biomarker method is not sufficient to predict all the variations of a heterogeneous disease like ovarian cancer (Koutsogiannouli et al., 2013). Microarray experiments studies are producing massive quantities of gene expression and other functional genomic data which gives us a deeper systems-level insight of a disease. These high-throughput genomic data were analyzed using various computational approaches to understand the mechanism of disease development for improved clinical diagnosis and therapeutics. Particularly co-expression network analysis is a powerful approach for finding the candidate genes based on similar expression pattern across the samples and these genes might be involved in similar biological pathways (Lee et al., 2004; van Dam et al., 2017; Sun et al., 2017; Han et al., 2018).

Extensive efforts have been made to identify diagnostic and prognostic genes in serous type of EOC using transcriptomic data (Ouellet et al., 2006; Hong et al., 2011; Sun et al., 2017; Han et al., 2018; Zhou et al., 2018). On the other hand, a limited number of studies were focused on the other subtypes of EOC where the comparative gene expression profiling among the different histologic subtypes of EOC were performed. Madore et al. (2010) performed a study to define the molecular characteristics of serous and endometrioid carcinomas to address the problems with the current histopathological classification methods. In another study, Pamula-Pilat et al. (2014) analyzed gene expression profiles in three histologic subtypes of EOC namely serous, endometrioid and clear cell carcinomas to identify molecular differences among the subtypes. Their study reported major molecular differences were observed between clear cell and serous carcinomas (1070 genes), and minor between endometrioid and serous carcinomas (81 genes). Recently, Cao and Wang (2018) reported a set of 1,265 differentially expressed genes (DEGs) were common between clear cell and serous carcinomas. In another recent work reported 13 hub genes those were functionally interacted among the 4 major subtypes of EOC and played crucial role in ovarian cancer progression (Singh and Som, 2018).

In this study, differential expression analysis was performed for four different subtypes of EOC along with healthy ovarian samples. Functional and pathway enrichment analyses of the identified DEGs were performed that showed the involvement of the DEGs in cancer related pathways. Further, a gene co-expression network of the DEGs in serous, endometrioid, mucinous and clear cell carcinomas were constructed separately with a query-based method followed by module analysis and eventually 15 common candidate genes were identified in all four subtypes. Further downstream analysis (such as survival analysis) of the candidate genes revealed a total of nine genes with significant prognostic value are proposed as the potential biomarker genes.

## 2. MATERIALS AND METHODS

### 2.1. Data retrieval

Gene expression datasets GSE6008 and GSE44104 were retrieved from the Gene Expression Omnibus (https://www.ncbi.nlm.nih.gov/geo/). The expression dataset GSE6008 incorporated 4 normal ovarian samples, 37 endometrioid, 41 serous, 13 mucinous, and 8 clear cell carcinomas samples (Hendrix et al., 2006). The dataset GSE44104 included 12 endometroid, 28 serous, 9 mucinous and 12 clear cell carcinoma samples (Wu et al., 2014).

### 2.2. Data processing

Raw microarray data contains different sources of noise such as due to the experimental procedure, efficacy and efficiency of the probe, noisy signal due to labelling efficiency, between slide variation and other factors. Hence, it is necessary to pre-process the raw data before the expression measures that can be used for further analysis. The raw data were standardized and transformed into expression value using the affy package of Bioconductor (Gautier et al., 2004). Pre-processing of the raw data was done separately. The robust multi-array average (RMA) algorithm was used for pre-processing of microarray data that included background signal correction, data normalization and probe summarization (Irizarry et al. 2003). The quality of the data was assessed through the boxplot.

### 2.3. Differential expression analysis

Linear models for microarray data (Limma) package was used to screen the DEGs between different subtypes of EOC by comparing it with the normal ovarian surface epithelium samples (Ritchie et al, 2015). Further, false-discovery rate arising due to multiple hypothesis testing were minimized through Benjamini-Hochberg’s method (Quackenbush, 2001). Both the criteria of absolute logarithmic fold change (|log_2_fc|)>1 and adjusted p-value (p_adj_)<0.05 were taken into account for the identification of DEGs (Yang et al., 2004).

### 2.4. Gene ontology and pathway enrichment analysis of DEGs

Gene ontology (GO) and KEGG pathway enrichment analysis of the DEGs were conducted using DAVID online tool that identify altered biological functions and pathways (Sherman al., 2007). GO terms consisted of three categories, including biological process (BP), molecular function (MF), and cellular component (CC). Significance testing of the matched terms were performed and a p-value<0.05 was considered to select the significantly enriched terms.

### 2.5. Co-expression network construction

Co-expression networks of the DEGs in four subtypes of EOCs were constructed by using the STRING database with the confidence score greater than or equal to 0.7 and visualized in Cytoscape (Shannon et al., 2003). The MCODE clustering algorithm was used which detects densely connected regions (module) in the co-expression networks by using following parameters: degree cutoff=2, node score cutoff=0.2, K-core=0.2 and maximum depth=100 (Bader et al., 2003). Further, we identified the genes and their interactions that were among the largest cluster of each co-expression network.

### 2.6. Survival analysis

Overall survival analysis of the genes that were shared among largest cluster of each subtype of EOC were performed by using Kaplan–Meier plotter online tool. Kaplan–Meier survival curves were used to assess the effect of genes on the overall survival of the ovarian cancer patients (Nagy et al., 2018). Significant differences in survival curves were assessed using the log-rank test (p-value<0.05) and hazard ratio (HR) with a 95% confidence interval.

### 2.7. Expression validation of core genes using TCGA and GTEx data

To further analyse the expression pattern of the genes that have significant prognostic value in the EOCs. We used GSE63885 microarray data and Gene Expression Profiling Interactive Analysis (GEPIA): a web-based server for the expression profiling of key genes in the tumor and normal samples (Tang et., 2017) using a threshold of |logfc|>1 and p_adj_<0.01. GEPIA perform differential expression analysis, profile plotting, correlation analysis, patient survival analysis, similar gene detection and dimensionality reduction analysis using RNA sequencing data from TCGA and GTEx projects (Tang et al., 2017)

## 3. RESULTS

### 3.1. Identification of DEGs in the four subtypes of EOC

Differential expression analysis was carried out to find the genes that are differentially expressed between cancerous and normal conditions. Here, we identified DEGs in the selected subtypes of EOC by using limma package for each dataset separately. Genes with p_adj_<0.05 and |log_2_fc|>1 were considered significantly differentially expressed. The proportion of genes found to be differentially expressed in each dataset is represented by the volcano plot shown in Figure 1. Table 1 summarizes the total number of upregulated and downregulated genes in serous, endometrioid, mucinous, and clear cell carcinomas. For the further downstream analyses, we considered the common DEGs in both datasets for each subtype. In the case of serous carcinoma, a total of 705 (upregulated=339 and downregulated=366) were common. Similarly, 649 (upregulated=290 and downregulated=359) genes in endometrioid, 835 (upregulated=408 and downregulated=427) genes in mucinous, and 689 (upregulated=318 and downregulated=371) genes were common in both datasets. Among all subtypes a total of 300 DEGs were common (Figure 2).

**Table 1:** Differentially expressed genes in different subtypes of epithelial ovarian cancer

| Data | Cancer type | Upregulated genes | Downregulated genes | Total |
| --- | --- | --- | --- | --- |
| GSE6008 | Serous carcinoma | 482 | 861 | 1343 |
|  | Endometrioid carcinoma | 431 | 795 | 1226 |
|  | Mucinous carcinoma | 2561 | 1709 | 4270 |
|  | Clear cell carcinoma | 613 | 1155 | 1768 |
| GSE44104 | Serous carcinoma | 1749 | 1214 | 2963 |
|  | Endometrioid carcinoma | 2005 | 795 | 3391 |
|  | Mucinous carcinoma | 2561 | 1709 | 4270 |
|  | Clear cell carcinoma | 1751 | 1260 | 3011 |

**Figure 1:**
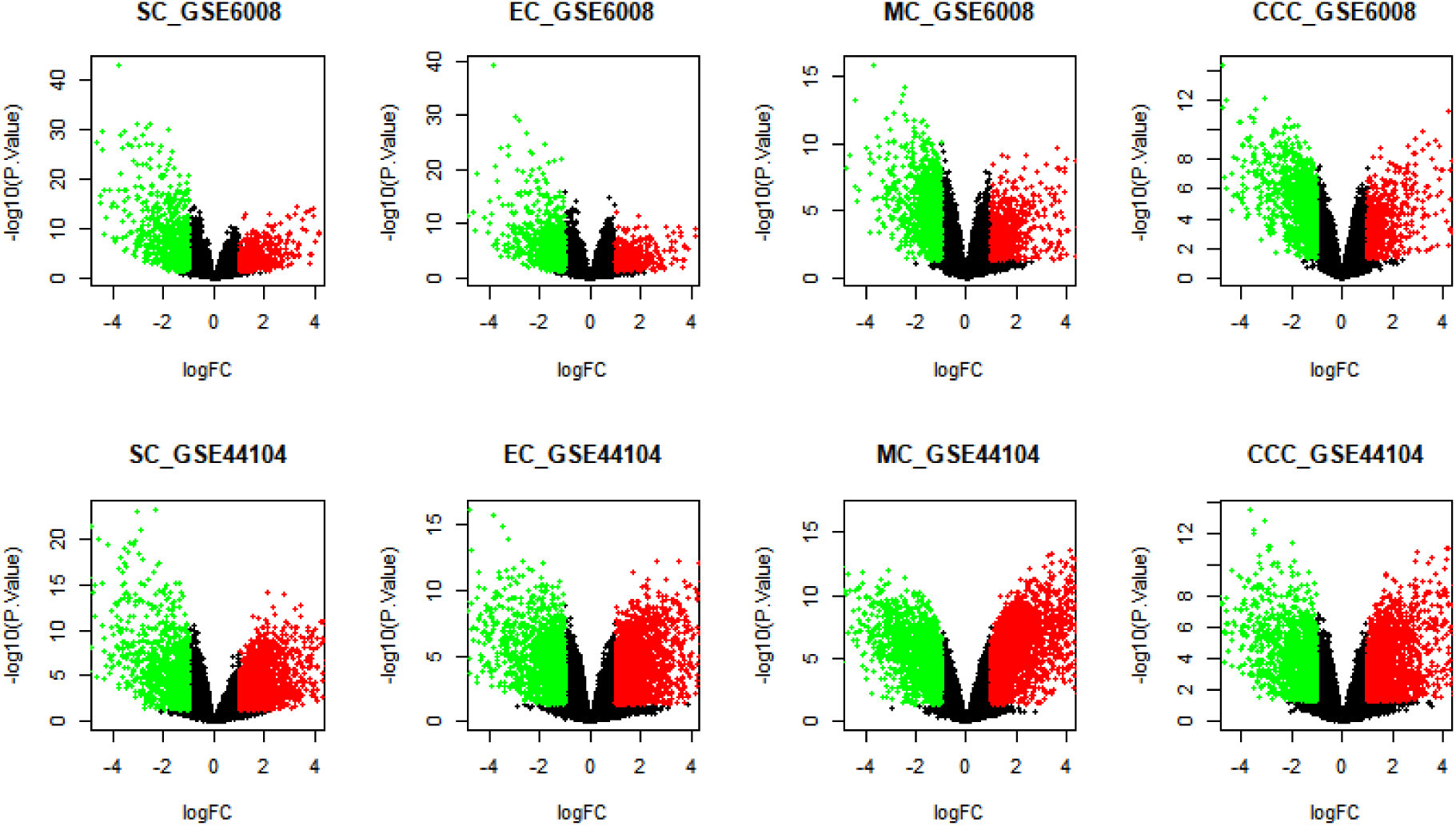
Volcano plot represents the proportion of genes found to be differentially expressed in each subtype of EOC within two datasets. The X-axis represents the log2 transformed of fold change ratios and Y-axis is the log10 transformed adjusted p-value. Green dots; down regulated DEGs, red dot; Up regulated DEGs based on |logfc|>1 and p<0.05.

**Figure 2:**
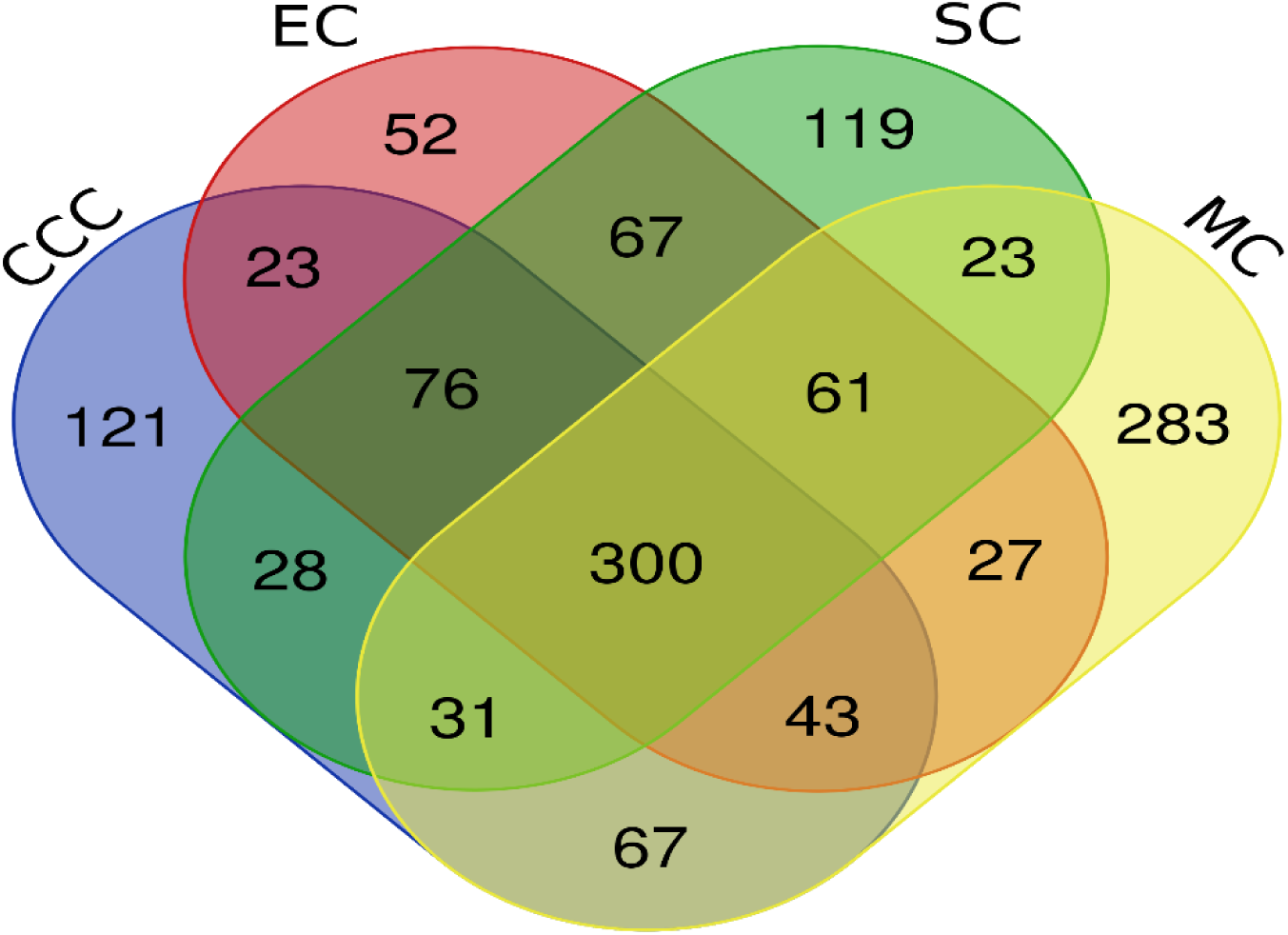
Venn diagram shows common DEGs among serous carcinoma (SC), endometrioid carcinoma (EC), mucinous carcinoma (MC), and clear cell carcinoma (CCC).

### 3.2. Function and pathway enrichment analysis of DEGs

To reveal the biological significance of DEGs in the progression of EOCs, GO and pathway enrichment analyses were performed using the DAVID online analysis tool with a p-value<0.05. Animal organ/cell development was the most significant biological process in the selected cancer subtypes. Similarly, protein kinase binding was the most significant molecular function and extracellular exome was most significant cellular component term for all four subtypes (Figure 3A-D).

**Figure 3A:**
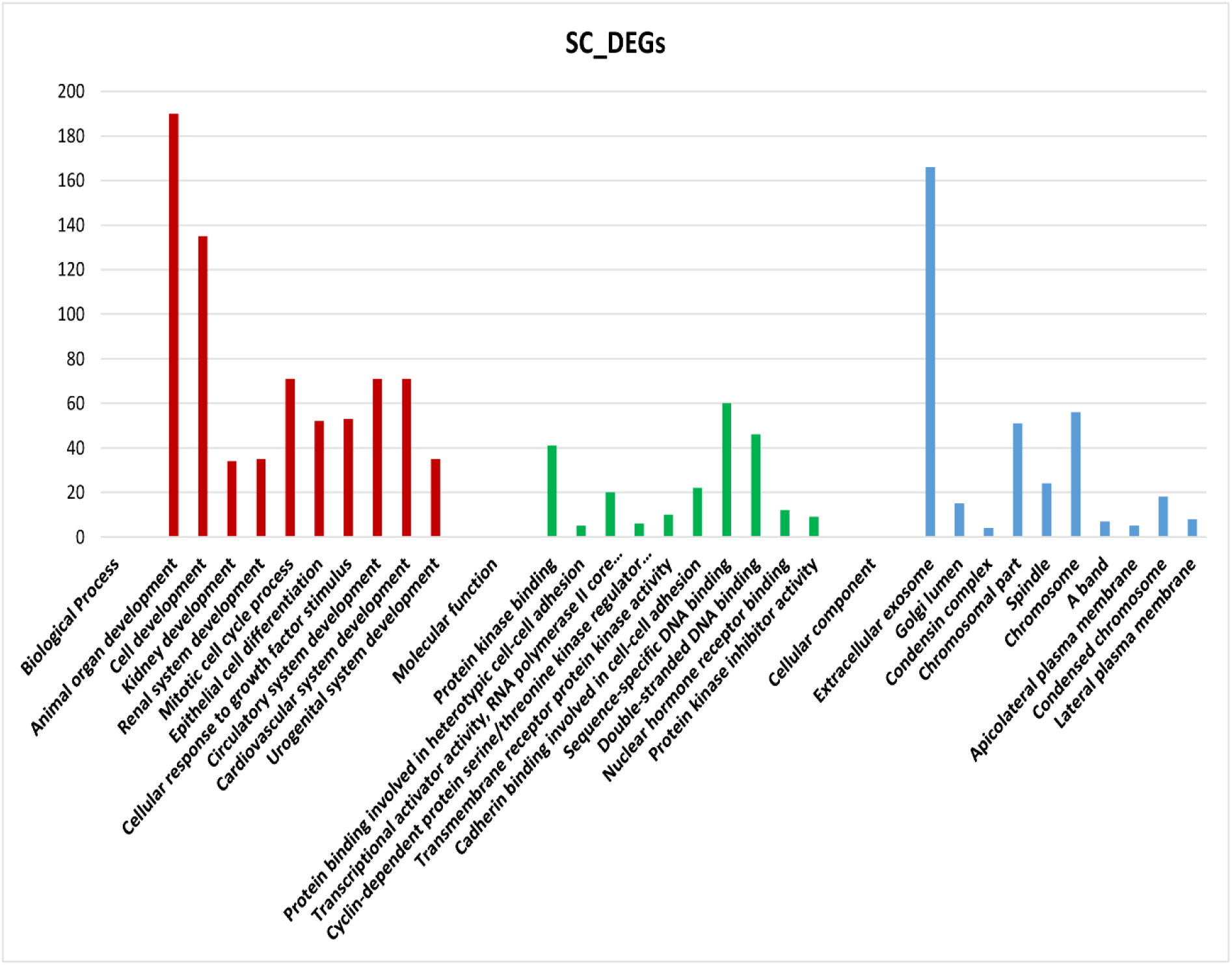
Represents the top 10 significantly enriched terms of the DEGs in serous carcinoma.

**Figure 3B:**
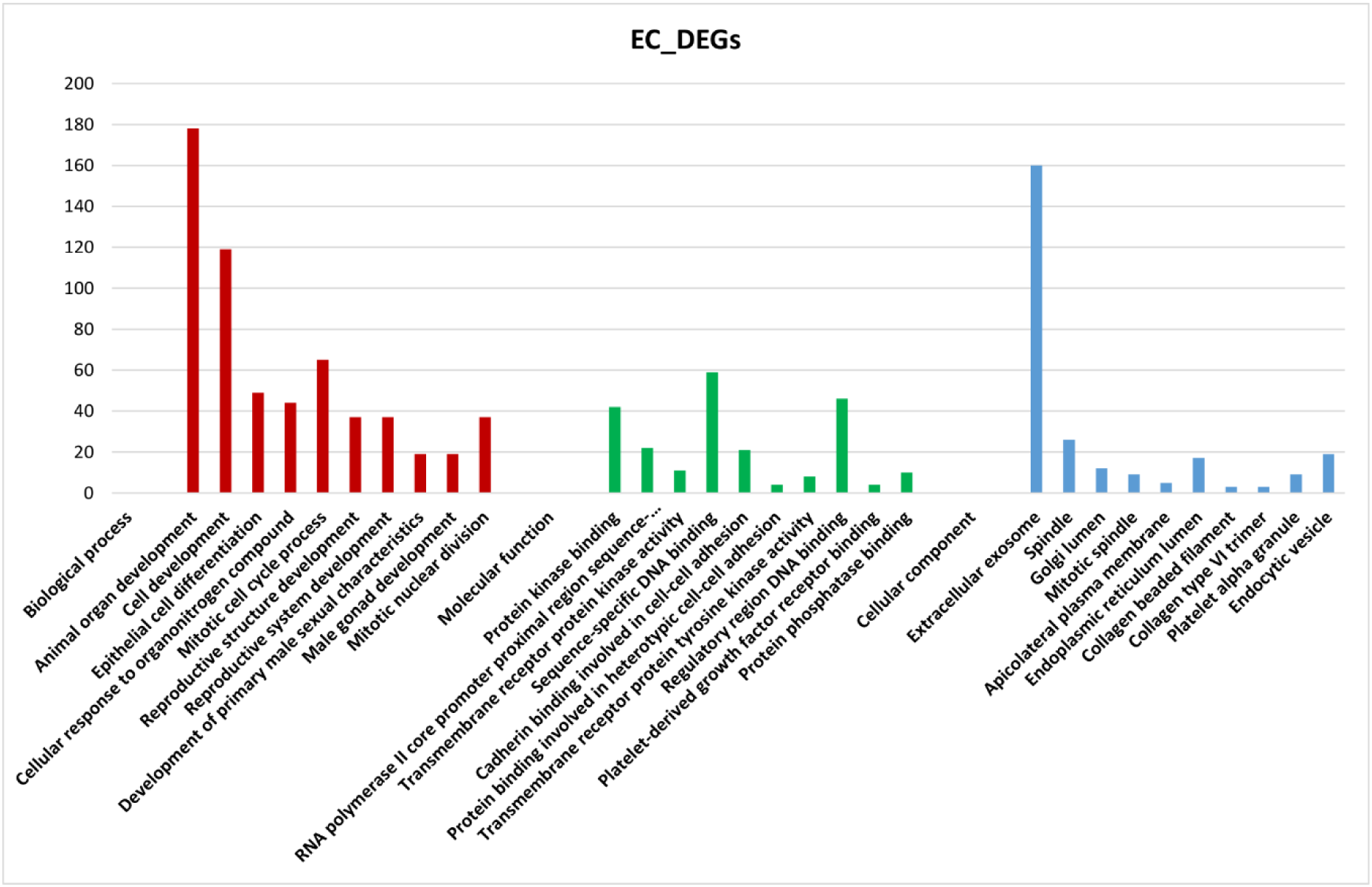
Represents the top 10 significantly enriched terms of the DEGs in endometroid carcinoma.

**Figure 3C:**
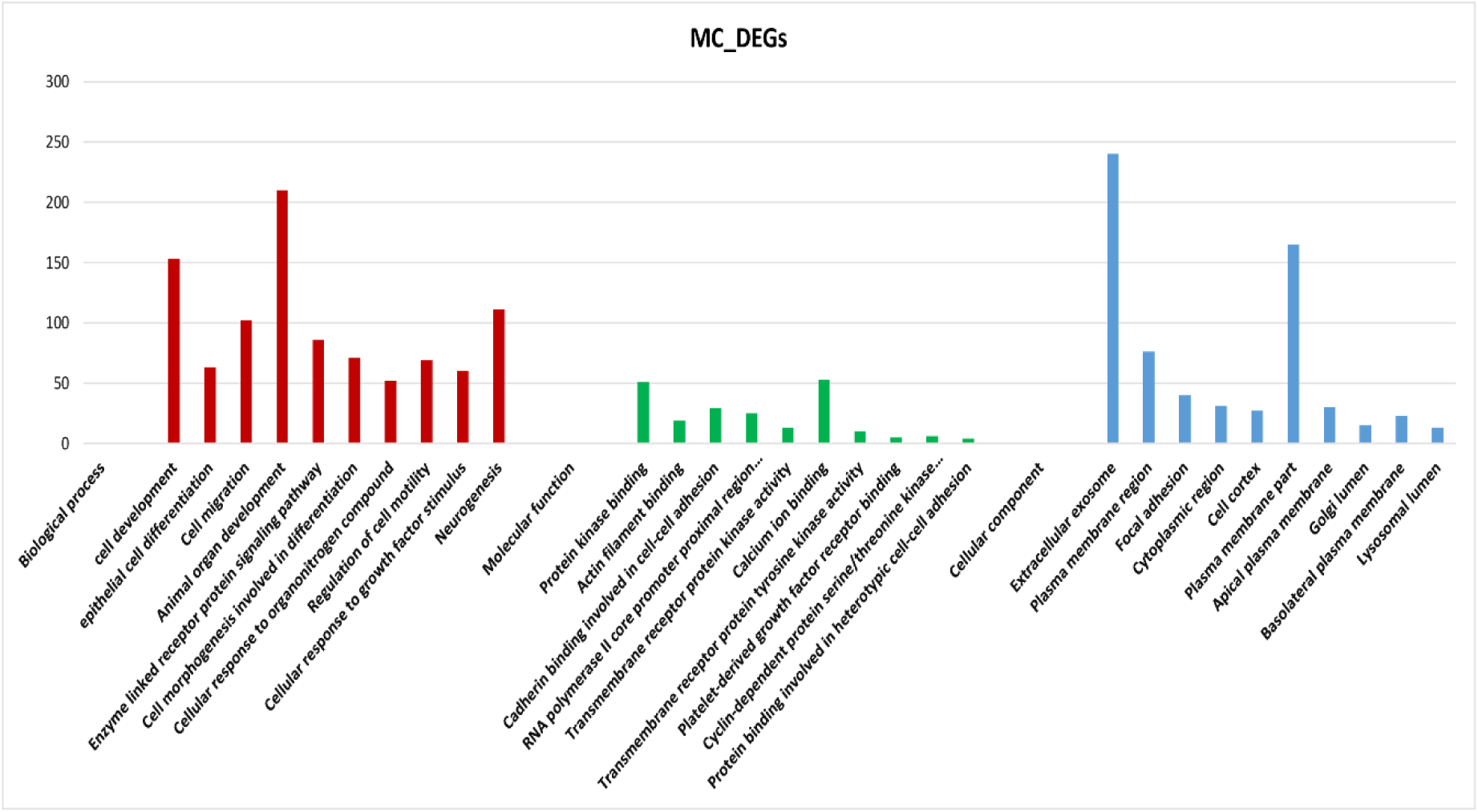
Represents the top 10 significantly enriched terms of the DEGs in mucinous carcinoma.

**Figure 3D:**
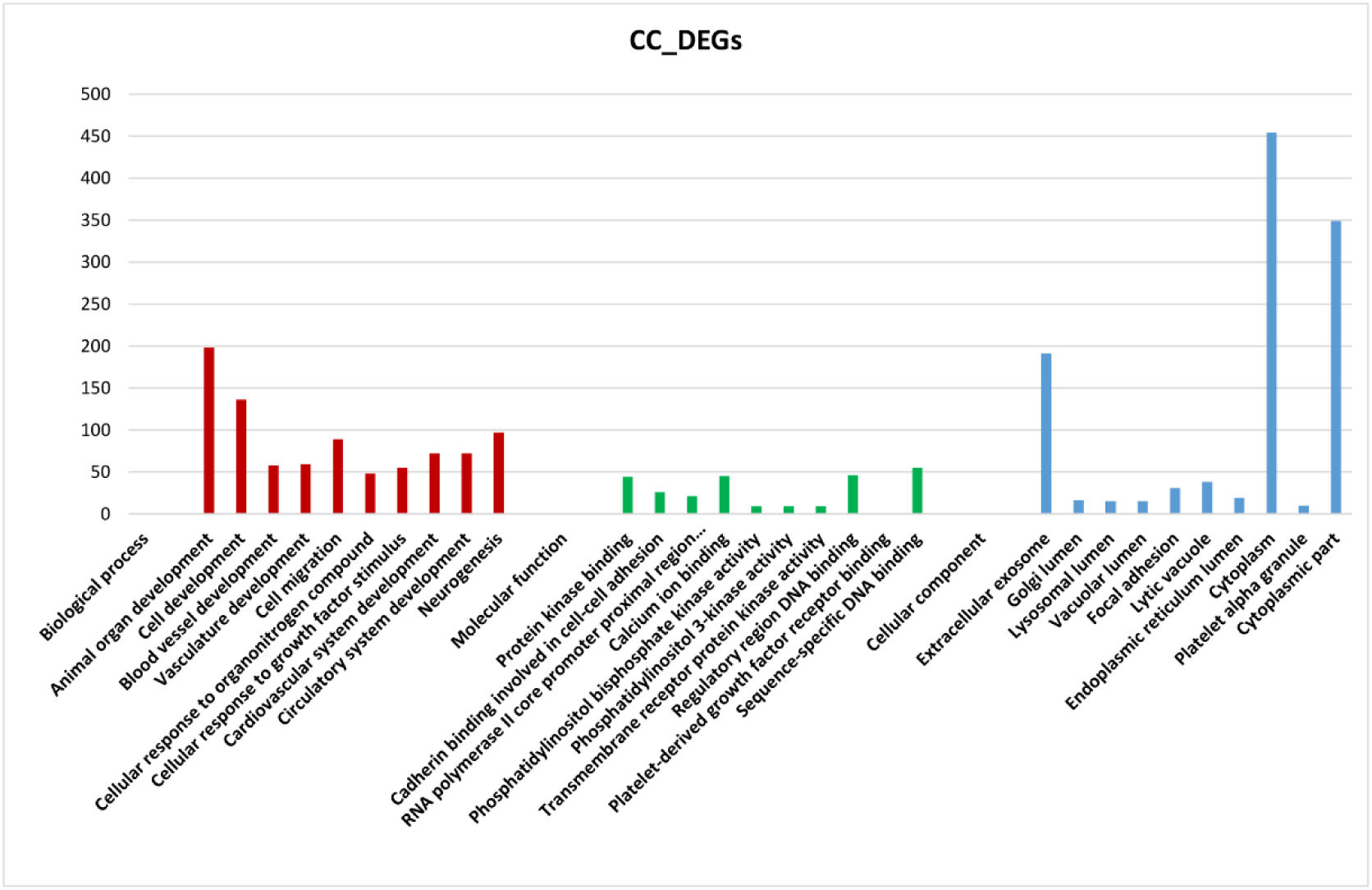
Represents the top 10 significantly enriched terms of the DEGs in clear cell carcinoma.

Pathway enrichment analysis of the DEGs in the selected cancer subtypes were carried out to identify pathways that are abnormally regulated and common across these subtypes. It had been identified that significant pathways are associated with the DEGs in serous, endometrioid, mucinous, and clear cell carcinomas with the p-value<0.05. Pathways in cancer, PI3K-AKT signaling pathway, RAP1 signaling pathway, cell cycle, Cell adhesion molecules, and proteoglycans in cancer are common pathways that were shared among the subtypes of EOC (Table 2).

**Table 2:** Common pathways involved in serous, endometroid, mucinous and clear cell carcinomas

| Term | Description |
| --- | --- |
| hsa04151 | PI3K-Akt signaling pathway |
| hsa05200 | Pathways in cancer |
| hsa04514 | Cell adhesion molecules |
| hsa04015 | Rap1 signaling pathway |
| hsa04110 | Cell cycle |
| hsa05205 | Proteoglycans in cancer |

### 3.3. Co-expression network of DEGs in each subtype of EOC

Co-expression networks and their modules directly reflect sample-specific interactions among the DEGs. Finding the common molecular interaction in different subtypes of EOC is important to understand diseases mechanism. We built co-expression network of DEGs in the four subtypes of EOC separately with the confidence score>0.7, a total of 73 DEGs in serous carcinoma formed co-expression network. Similarly, 76 DEGs in endometrioid carcinoma, 49 DEGs in mucinous carcinoma, and 57 genes in clear cell carcinoma were formed co-expression networks (Supplementary File 1). Further co-expression module of each cancer subtypes was identified using MCODE clustering algorithm. The largest module in serous carcinoma has 41 genes with 773 interactions, endometrioid carcinoma has 39 genes with 669 interactions, mucinous has 20 genes with 171 interactions, and clear cell carcinoma has 23 genes with 223 interactions (Figure 4A-D). Finally, by merging the largest modules of different subtypes of EOC, it had been derived a network of 15 genes (candidate genes) having 101 interactions that reflects the common molecular mechanisms in the four subtypes of EOC (Figure 5). Details of the candidate genes are given in Table 3.

**Table 3:** List of the identified common candidate genes involved in the progression of serous, endometroid, mucinous and clear cell carcinomas

| Gene symbol | Ensembl id | Description |
| --- | --- | --- |
| HJURP | ENSG00000123485 | Holliday junction recognition protein |
| KIF4A | ENSG00000090889 | Kinesin family member 4A |
| UBE2C | ENSG00000175063 | Ubiquitin conjugating enzyme E2C |
| CEP55 | ENSG00000138180 | Centrosomal protein 55 |
| PTTG1 | ENSG00000164611 | Pituitary tumor-transforming gene-1 |
| ASPM | ENSG00000066279 | Abnormal spindle microtubule assembly |
| CENPM | ENSG00000100162 | Centromere protein M |
| RACGAP1 | ENSG00000161800 | Rac GTPase activating protein 1 |
| TPX2 | ENSG00000088325 | Targeting protein for Xenopus kinesin like protein 2 |
| ZWINT | ENSG0000122952 | ZW10 interacting kinetochore protein |
| CDCA8 | ENSG00000134690 | Cell division cycle associated 8 |
| MELK | ENSG00000165304 | Maternal embryonic leucine zipper kinase |
| CDC20 | ENSG00000117399 | Cell division cycle 20 |
| TOP2A | ENSG00000131747 | DNA topoisomerase II alpha |
| HMMR | ENSG00000072571 | Hyaluronan mediated motility receptor |

**Figure 4A:**
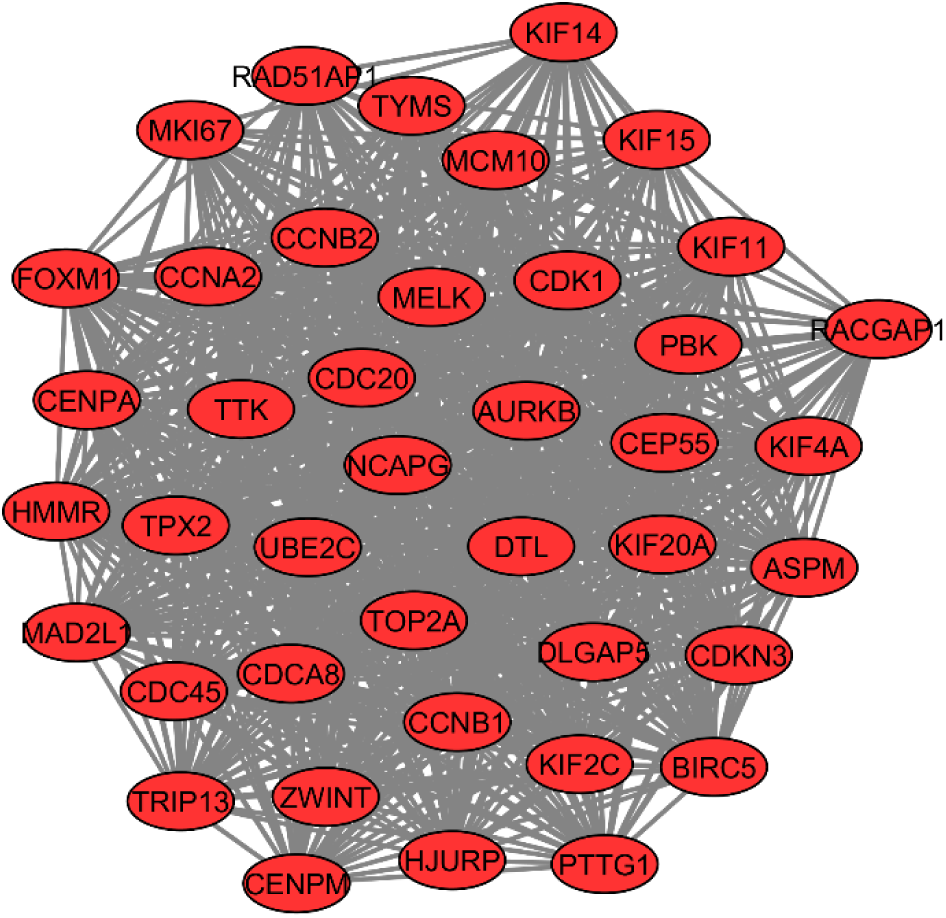
Shows the largest co-expression modules in serous carcinoma. Red color nodes represent upregulated genes.

**Figure 4B:**
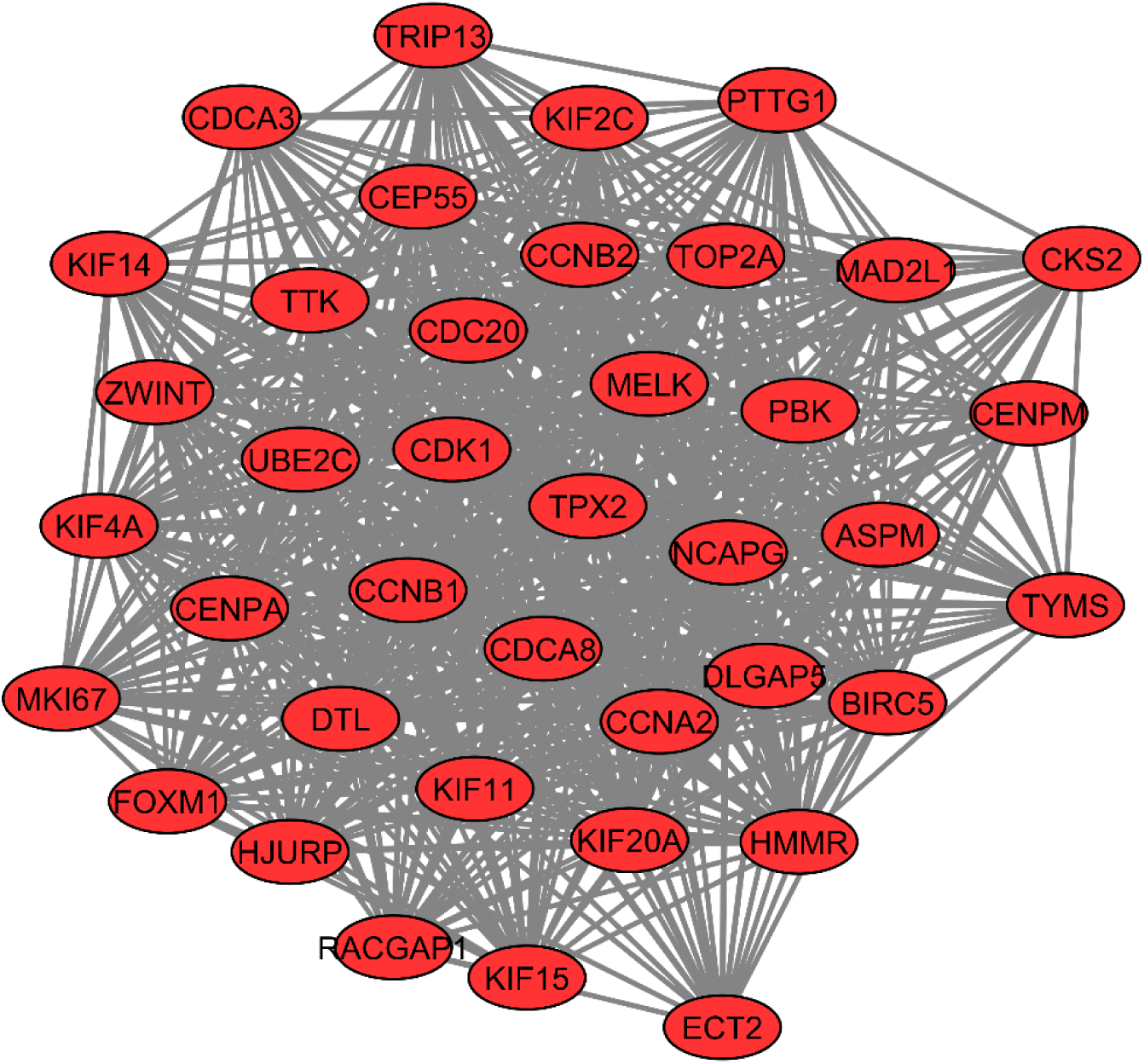
Shows the largest co-expression module in endometrioid carcinoma. Red color nodes represent upregulated genes.

**Figure 4C:**
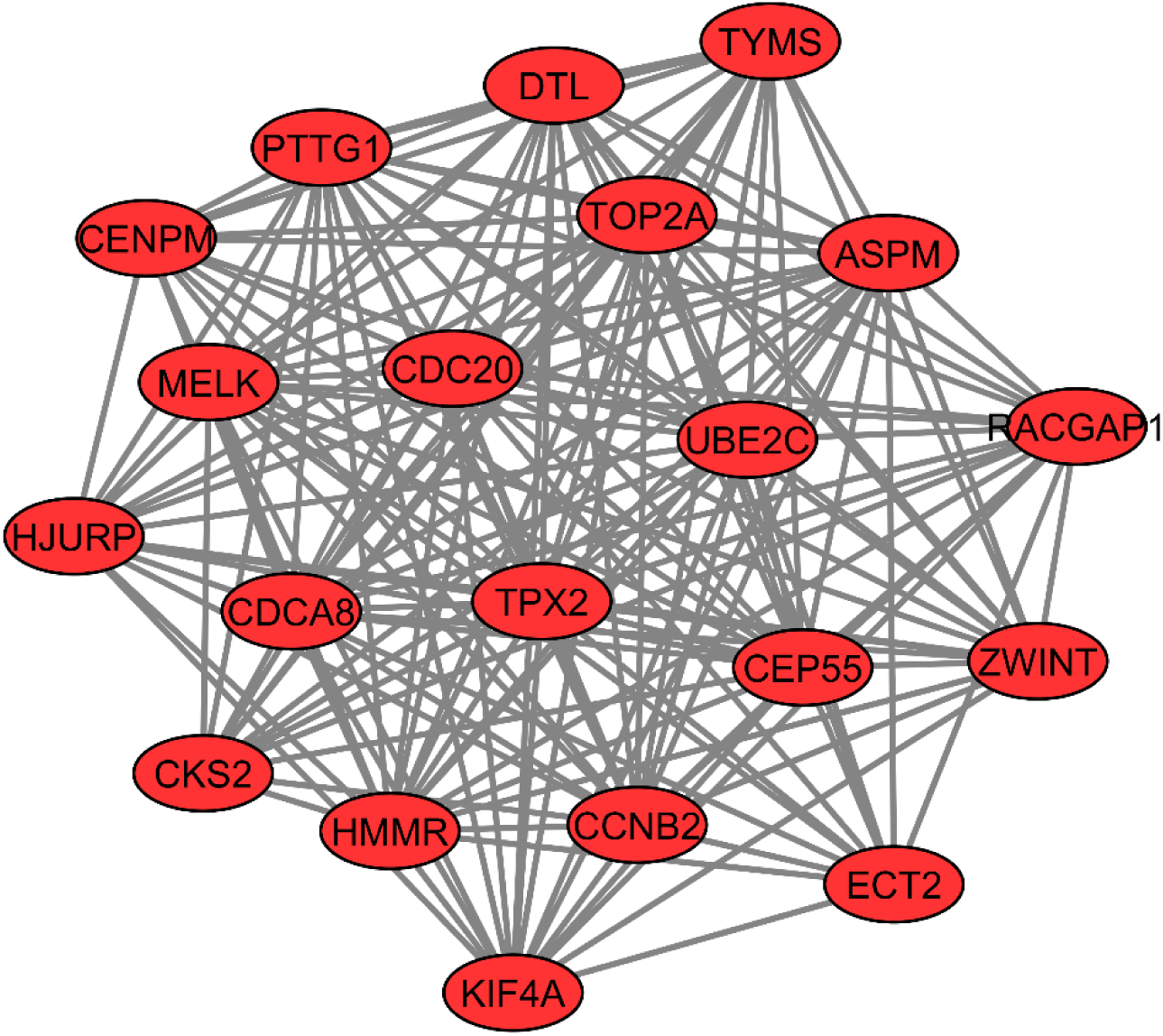
Shows the largest co-expression module in mucinous carcinoma. Red color nodes represent upregulated genes.

**Figure 4D:**
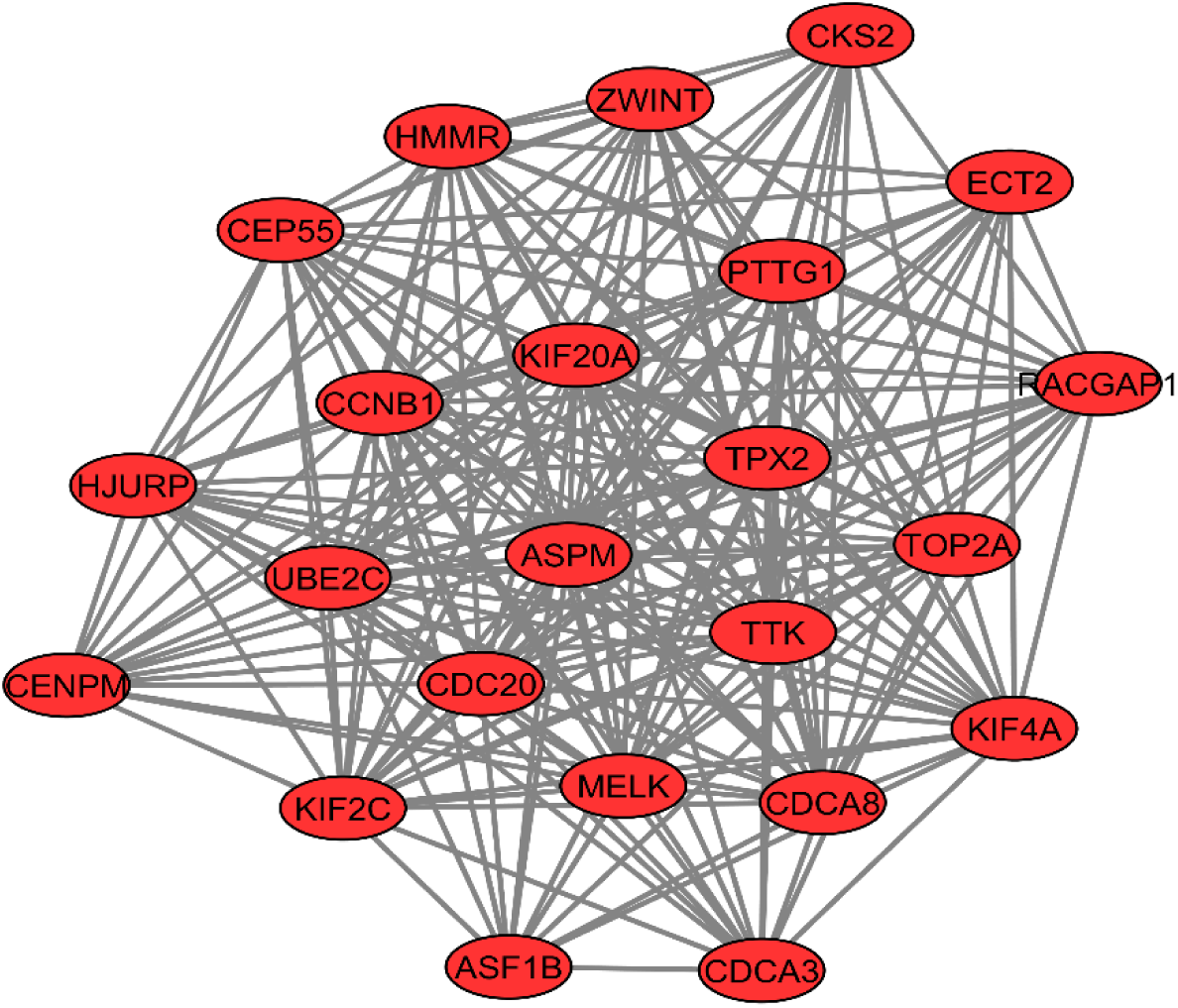
Shows the largest co-expression module in clear cell carcinoma. Red color nodes represent upregulated genes.

**Figure 5:**
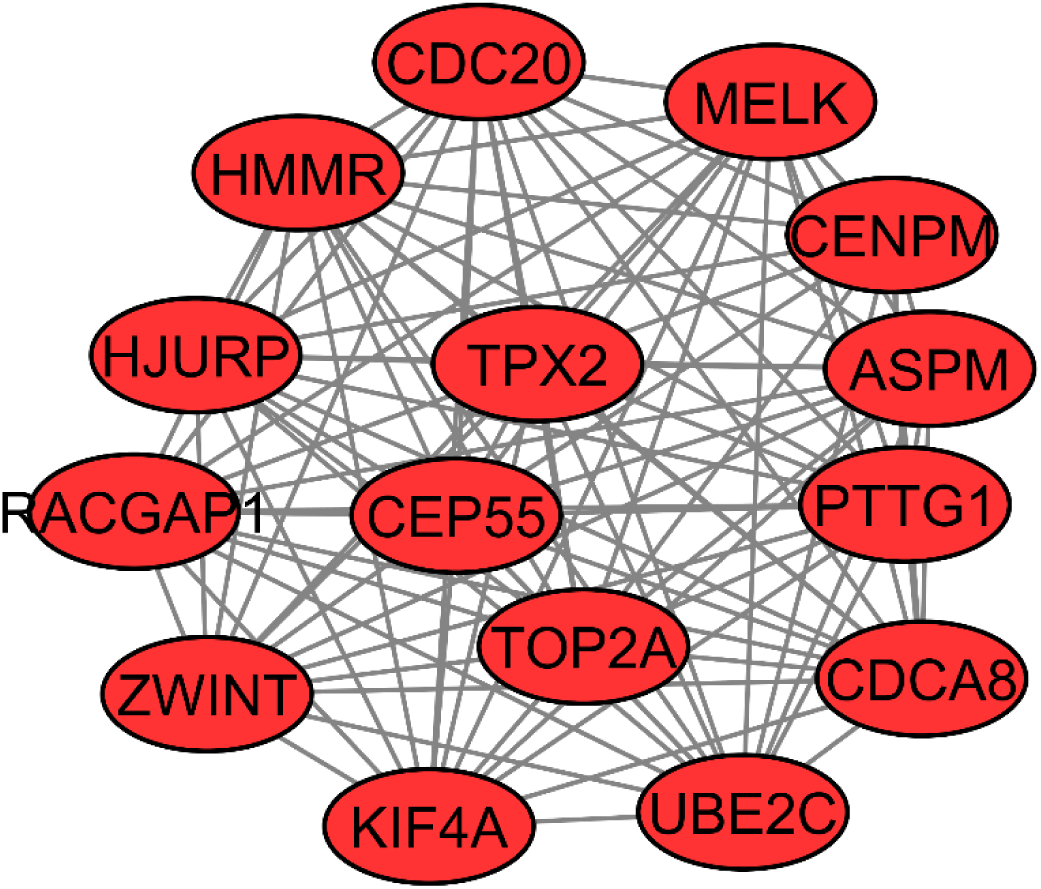
Network of 15 co-expressed genes that were common among the largest module of each subtype of EOC.

### 3.4. Overall survival analysis of the candidate genes

Kaplan-Meier plotter was used to analyze the effect of identified common candidate genes on the survival of patients having epithelial ovarian cancer. It had been identified that out of 15 candidate genes nine genes namely ASPM, CDCA8, CENPM, CEP55, HMMR, RACGAP1, TPX2, UBE2C, and ZWINT have significant prognostic value and high mRNA expression of these genes were associated with a poor overall survival of EOC patients. Hence, these nine genes are reported as potential biomarkers (Figure 6). Details of the biomarker genes are given in Table 4.

**Table 4:**
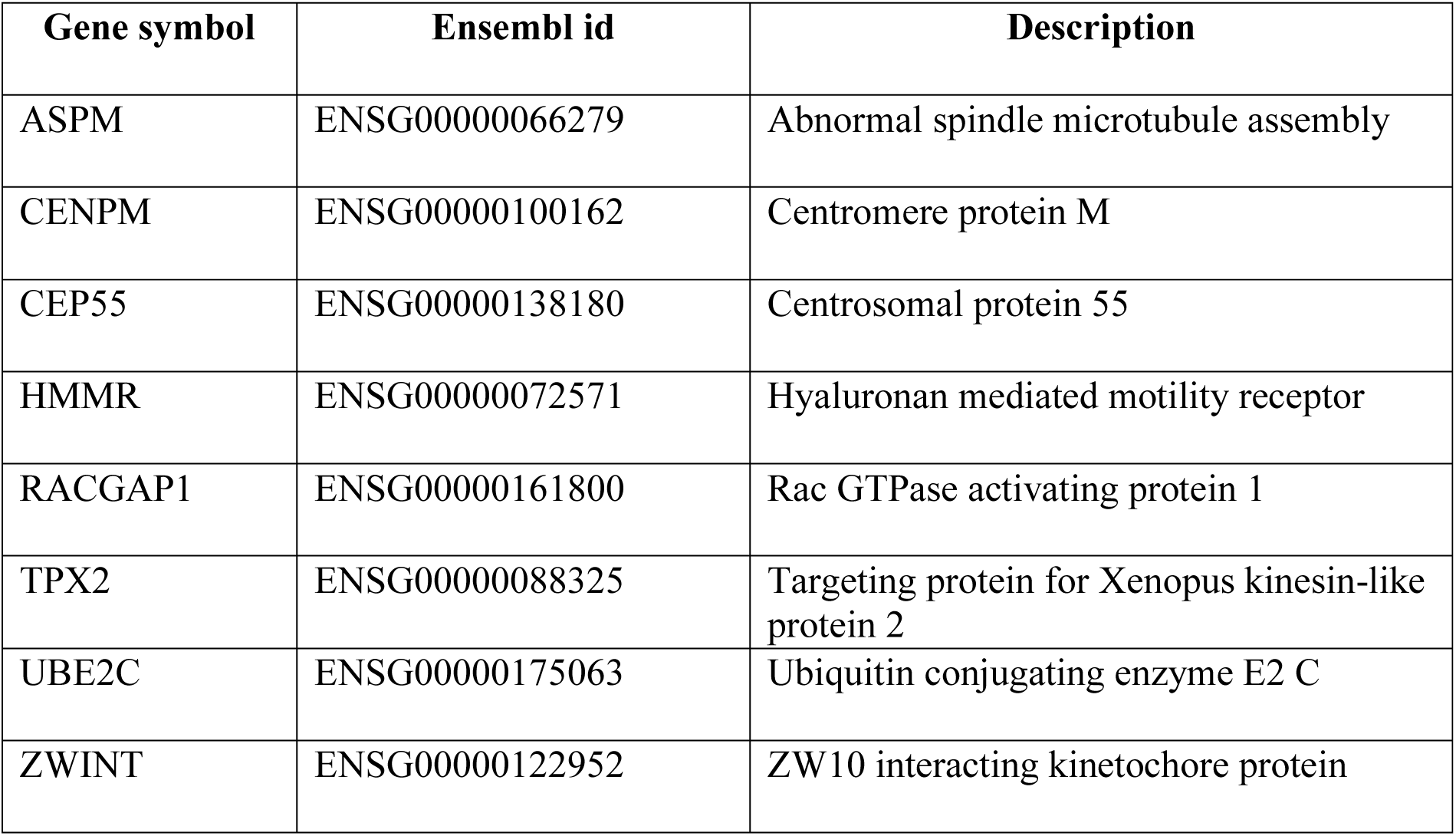
List of the identified common biomarker genes involved in the progression of serous, endometroid, mucinous and clear cell carcinomas

| Gene symbol | Ensembl id | Description |
| --- | --- | --- |
| ASPM | ENSG00000066279 | Abnormal spindle microtubule assembly |
| CENPM | ENSG00000100162 | Centromere protein M |
| CEP55 | ENSG00000138180 | Centrosomal protein 55 |
| HMMR | ENSG00000072571 | Hyaluronan mediated motility receptor |
| RACGAP1 | ENSG00000161800 | Rac GTPase activating protein 1 |
| TPX2 | ENSG00000088325 | Targeting protein for Xenopus kinesin-like protein 2 |
| UBE2C | ENSG00000175063 | Ubiquitin conjugating enzyme E2 C |
| ZWINT | ENSG00000122952 | ZW10 interacting kinetochore protein |

**Figure 6:**
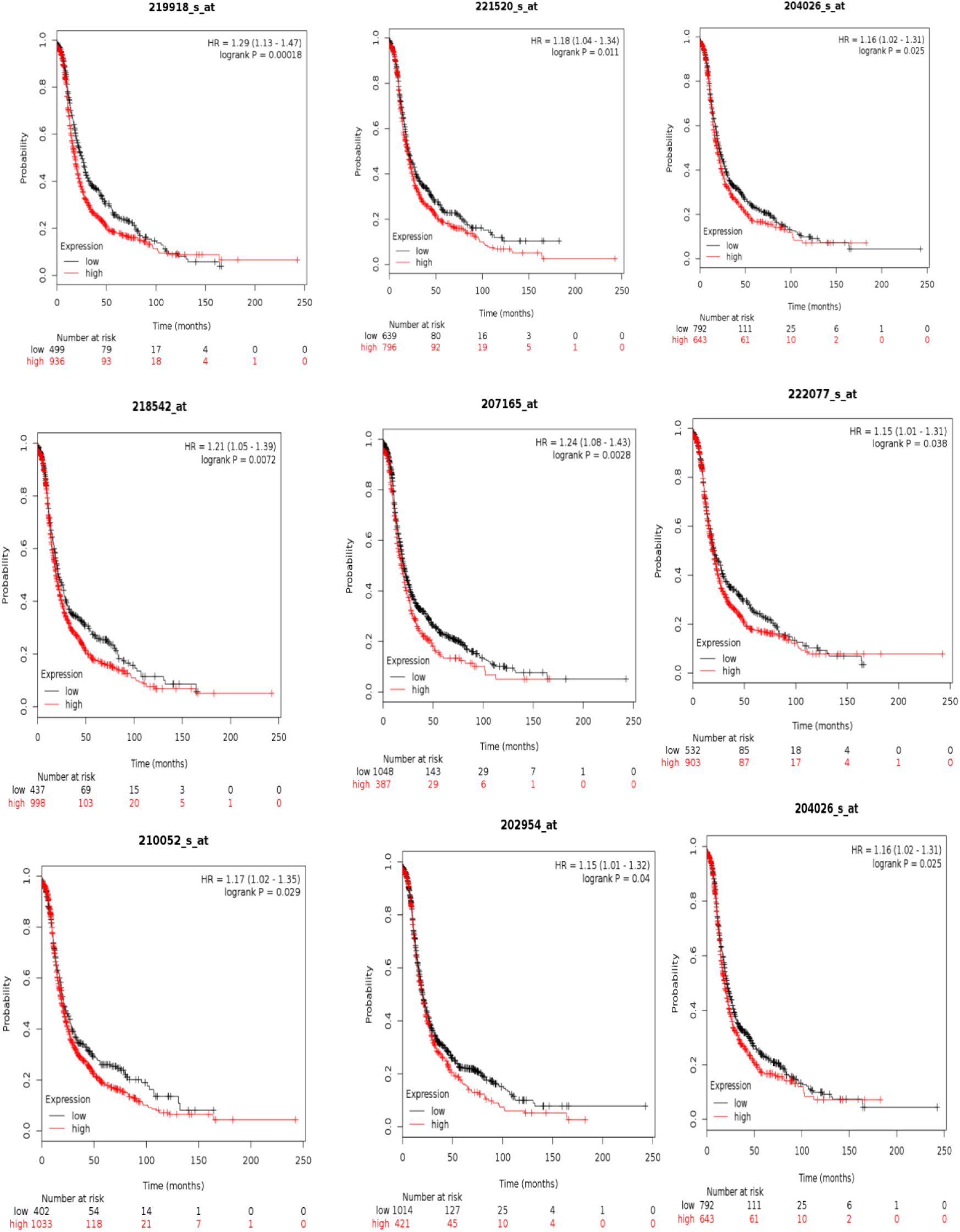
Kaplan-Meier survival curves. High expression of the biomarker genes was associated with overall poor survival of EOC patients. 219918_at; ASPM, 221520_at; CDCA8, 204026_at; ZWINT, 218542_at; CEP55, 207165_at; HMMR, 222077_s_at; RACGAP1, 210052_at; TPX2, 202954_at; UBE2C, 218741_at; CENPM.

### 3.5. Expression validation of core genes using TCGA and GTEx data

To further analyse the expression pattern of the candidate genes that have significant prognostic value in the EOCs at the criteria of |logfc|>1 and p_adj_<0.01, we used GSE63885 microarray data and GEPIA web-based server. It had been identified that these genes were overexpressed in different histologic subtypes of EOC as compared to normal ovarian samples (Figure 7).

**Figure 7:**
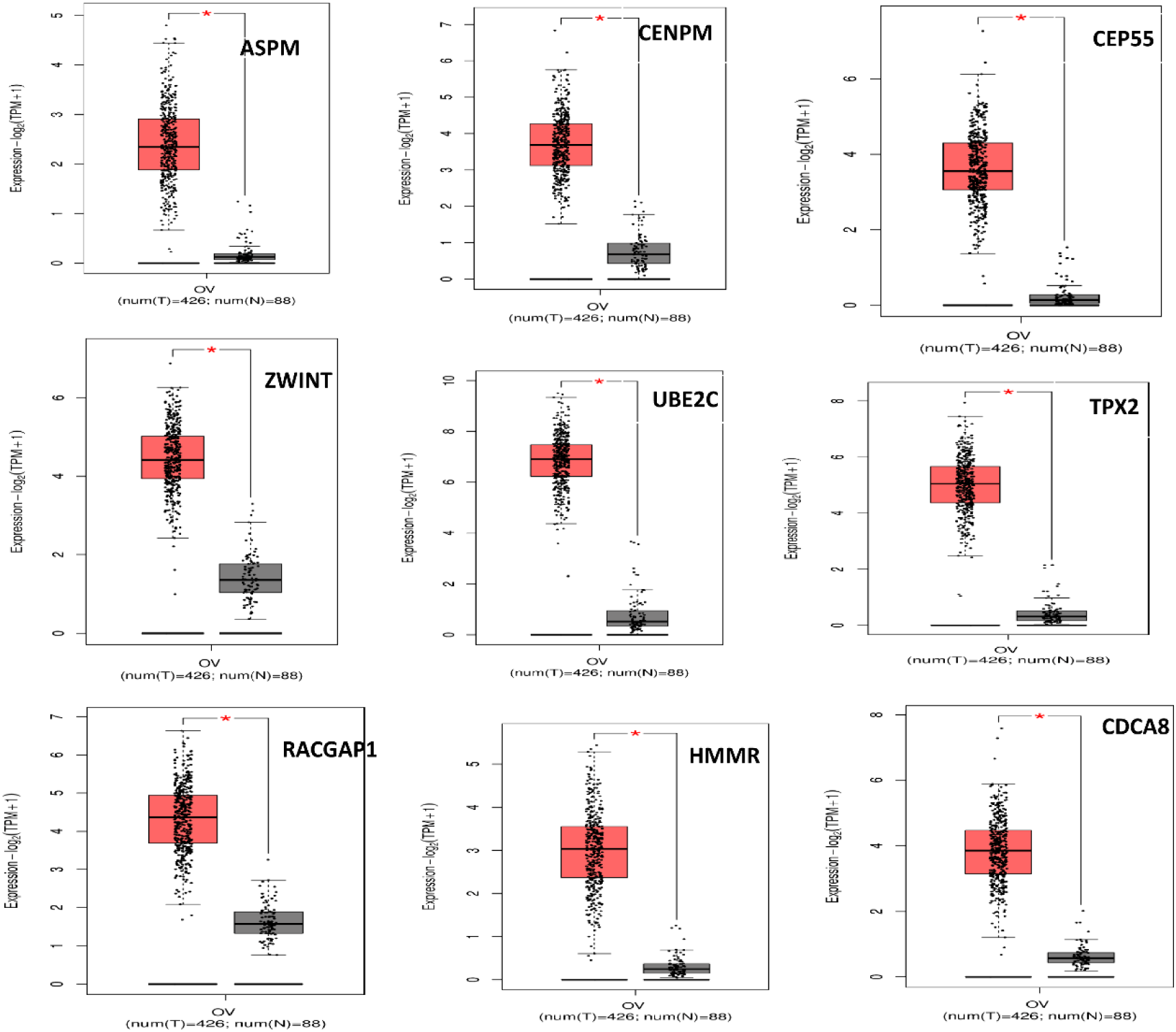
Box plots represent the differential expression of the biomarker genes that have significant prognostic value. Red boxes indicate tumor samples and grey boxes indicate normal samples.

Furthermore, GO analysis of the 15 candidate genes showed their significantly involvement in the cell cycle, cell proliferation and cytoskeleton organisation process.

## 4. DISCUSSION

Epithelial ovarian cancer is one of the most lethal gynecological cancer worldwide due to heterogeneity, delayed diagnosis as well as recurrence and drug resistance. Thus, the development of new potential biomarker is required that might enable better treatment of EOC.

In the present study we identified DEGs in the four major subtypes of EOC. The GO and pathway enrichment analyses reveal common biological functions and pathways involved in carcinogenesis of the EOC, and further co-expression networks and module analysis reported 15 candidate genes that were common in the four subtypes of EOC and formed a core circuitry. Survival analysis of the candidate genes were showed that high expression of the nine genes namely ASPM, CDCA8, CENPM, CEP55, HMMR, RACGAP1, TPX2, UBE2C, and ZWINT were associated with epithelial ovarian carcinomas development that leads toward overall poor survival of epithelial ovarian cancer patients.

ASPM (Abnormal spindle-like microcephaly-associated protein) is essential for normal mitotic spindle function during cell division and also regulate neurogenesis (Kouprina et al., 2005). Growing evidence shows that abnormalities in ASPM expression are associated with numerous cancer types (Horvath et al., 2006; Lin et al., 2008; Pai et al., 2019). Previous studies have revealed that high expression of ASPM is often in ovarian and other human cancers links with poor clinical prognosis and early recurrence (Brüning-Richardson et al., 2011; Zhou et al., 2018; Xu et al., 2019).

CDCA8 (Cell division cycle associated 8) encodes a component of the chromosomal passenger complex (CPC). This complex is an essential regulator of mitosis and cell proliferation by regulating chromatin-induced microtubule stabilization and spindle formation. Transcription factor NF-Y is a positive regulator of CDCA8 transcription (Ruchaud et al., 2007). CDCA8 is a putative oncogene that is overexpressed in various human cancers including ovarian cancer and is required for cancer growth and progression (Zhou et al., 2018; Bu et al., 2019).

CENPM (Centromere protein M) also known as proliferation associated nuclear element plays a central role in assembly of kinetochore proteins, successful mitotic progression and chromosome alignment. Study of Basilico et al. (2014) demonstrated that CENP-M depletion also caused robust mitotic arrest, likely a consequence of spindle checkpoint activation caused by chromosome alignment defects. A recent study by Ding et al. (2019) identified novel promising biomarkers for the prognosis of cervical cancer. Tissue microarray and gene network analyses reported that CENPM is common therapeutic target of garlic and cisplatin in bladder cancer (Kim et al., 2018).

CEP55 (Centrosomal Protein 55), has long been recognized as scaffold proteins, regulating both the mitotic spindle and microtubule organization, and hence are critical for cell cycle progression. Its overexpression is associated with genomic instability, a hallmark of cancer (Xu et al., 2015). Study by Jeffery et al. (2016) reported that CEP55 is highly overexpressed in several human cancer and overexpression of CEP55 is associated with invasion of tumors. Overall, CEP55 is good prognostic and therapeutic target for the cancer treatment.

HMMR (hyaluronan mediated motility receptor) an oncogenic protein. HMMR protein involved in microtubule spindle assembly and also contributed to cell cycle regulation. On the extracellular surface, HMMR forms a complex with cluster differentiation 44 (CD44) and hyaluronan (HA) to activate cell signaling pathways that promote migration, invasion and cell proliferation (Misra et al., 2015). Further it had been reported that the overexpression of HMMR contribute to tumor progression, aggressive phenotype and poor prognosis in multiple cancer types (Maxwell et al., 2008).

RACGAP1 (Rac GTPase-activating protein 1), a central spindle complex plays very important role in controlling various cellular process including cytokinesis, transformation, invasive migration and metastasis (Zhao et al., 2005). Previous studies identified the overexpression of RACGAP1 in gastric, ovarian, colorectal and several other cancers, implied its role in promoting tumor progression (Imaoka et al., 2015; Saigusa et al., 2015, Wang et al., 2018).

TPX2 (Targeting protein for Xenopus kinesin-like protein 2) is one of the many microtubules associated proteins that plays a key role in microtubule assembly and growth during M phase by interacting directly or indirectly with several other proteins that regulate spindle assembly and function, including microtubule-binding proteins, motors and nucleation factors (Pérez de Castro et al., 2012). Several studies demonstrate that TPX2 is overexpressed in multiple cancer types and play important role in promoting tumorigenesis and metastasis (Liang et al., 2016; Zou et al., 2018). A recent study reported that TPX2/AURK signaling as potential target in genomic unstable cancer cell including ovarian cancer and breast cancer (Hsu et al., 2017; van Gijn et al., 2019).

UBE2C (Ubiquitin-conjugating enzyme E2C) protein is involved in the ubiquitination process that modifies the abnormal or short-lived proteins with ubiquitin and target toward degradation (van Wijk et al., 2010). A good number of studies showed high expression of UBE2C is associated with aggressive progression and poor outcomes of several cancer (Dastsooz et al., 2019).

ZWINT, as a part of kinetochore complex, is a protein interacting with ZW10 that plays essential role in kinetochore-microtubule attachment, spindle assembly check point function and ensures that chromosomes are divided equally between daughter cells (Wang et al., 2004). Recently, overexpression of ZWINT has been reported in human malignancies including prostate, ovarian, lung, and breast cancers, as well as lung cancer and hepatocellular carcinoma (Xu et al., 2016; Ying et al., 2018; Peng et al., 2019).

Thus, the nine biomarkers genes reported in this work along with the role of the genes explored by the other studies confirm that the genes namely ASPM, CDCA8, CENPM, CEP55, HMMR, RACGAP1, TPX2, UBE2C, and ZWINT involve in the pathogenesis of malignant tumors by affecting mitosis, and cell cycle process, which support our findings.

## 5. CONCLUSIONS

Differential gene expression analysis and network-based approach were used to analyse molecular profiles of four major subtypes of epithelial ovarian cancer. Our analysis revealed nine common potential biomarkers in four subtypes of epithelial ovarian cancer and their upregulation is associated with pathogenesis of epithelial malignant tumors and overall poor survival of the epithelial ovarian cancer patients by affecting mitosis, and cell cycle process. Hence, the reported biomarkers could be an interest of the experimentalists as potential therapeutic targets for the subtypes of epithelial ovarian cancer.

## ACKNOWLEDGMENTS

The authors would like to thank Mr. Arindam Ghosh, Ms. Priyanka Kumari and Mr. Amresh Sharma for various useful discussion and suggestions. AS gratefully acknowledge the DBT India for financial support.

## Supplementary File

**Supplementary File 1:** Cytoscape file consists of the co-expression networks of serous, endometrioid, mucinous and clear cell carcinoma.

